# Last in first out: SIV proviruses seeded later in infection are harbored in short-lived CD4^+^ T cells

**DOI:** 10.1101/2023.11.03.565539

**Authors:** Narmada Sambaturu, Emily J. Fray, Fengting Wu, Carolin Zitzmann, Francesco R. Simonetti, Dan H. Barouch, Janet D. Siliciano, Robert F. Siliciano, Ruy M. Ribeiro, Alan S. Perelson, Carmen Molina-París, Thomas Leitner

## Abstract

HIV can persist in a latent form as integrated DNA (provirus) in resting CD4^+^ T cells of infected individuals and as such is unaffected by antiretroviral therapy (ART). Despite being a major obstacle for eradication efforts, the genetic variation and timing of formation of this latent reservoir remains poorly understood. Previous studies on when virus is deposited in the latent reservoir have come to contradictory conclusions. To reexamine the genetic variation of HIV in CD4^+^ T cells during ART, we determined the divergence in envelope sequences collected from 10 SIV infected rhesus macaques. We found that the macaques displayed a biphasic decline of the viral divergence over time, where the first phase lasted for an average of 11.6 weeks (range 4-28 weeks). Motivated by recent observations that the HIV-infected CD4^+^ T cell population is composed of short- and long-lived subsets, we developed a model to study the divergence dynamics. We found that SIV in short-lived cells was on average more diverged, while long-lived cells harbored less diverged virus. This suggests that the long-lived cells harbor virus deposited starting earlier in infection and continuing throughout infection, while short-lived cells predominantly harbor more recent virus. As these cell populations decayed, the overall proviral divergence decline matched that observed in the empirical data. This model explains previous seemingly contradictory results on the timing of virus deposition into the latent reservoir, and should provide guidance for future eradication efforts.

**Significance statement:** HIV can persist in a latent reservoir unaffected by antiretroviral drugs. The genetic variation of this latent virus population is a major obstacle for eradication efforts, but also a clue to when HIV variants are deposited in the reservoirs. Unfortunately, previous studies assessing when the virus was deposited in latent reservoirs have come to contradictory conclusions. Here, we propose SIV proviral DNA exists in both short- and long-lived CD4^+^ T cells, and that these two cell subsets harbor different genetically diverged virus populations. Our model explains the contradictory findings and shows that when CD4^+^ T cells decay under effective drug treatment, which prevents virus replication, the resulting virus divergence decreases and recapitulates observed data. This knowledge should help in improving future eradication efforts.

---

Human immunodeficiency virus (HIV) has infected over 84 million people and caused an estimated 40.1 million deaths worldwide since the beginning of the epidemic in 1981 (1, 2). Antiretroviral therapy (ART) suppresses the replication of HIV, reducing viral load and prolonging life (1). However, even during suppressive ART, a latent reservoir of transcriptionally silent intact proviruses persists, integrated in the genomes of resting CD4^+^ T cells of the host (3–7). This provirus reservoir is capable of re-seeding infection if therapy is interrupted, making HIV infection as of yet incurable with ART alone (8, 9). Despite the importance of the latent reservoir, the impact of T cell dynamics on the timing of its formation remains largely unknown.

Attempts to identify when the latent reservoir is established have yielded seemingly contradictory results. Several studies have observed that even when ART is initiated early (within days of infection), viral rebound occurs upon treatment interruption (10– 12), suggesting early and continuous seeding of the latent reservoir. Other studies discovered that viral rebound sequences obtained during treatment interruption were genetically closest to the founder virus, also suggesting that the latent reservoir is established primarily near the start of infection (13–15). Still other studies found the latent reservoir to be genetically heterogeneous, recapitulating HIV’s within-host evolutionary history, suggesting that the latent reservoir is seeded throughout the course of infection (16–18). Comprehensive simulations of within-host sequence evolution that included a latent reservoir, immune selection, point mutations and recombination, demonstrated that empirical diversity and divergence trends were consistent with continuous deposition of HIV variants throughout natural infection (19). In contradiction, more recent studies have found that a majority of proviral DNA sequences, obtained several years after initiation of ART, clustered closer on a phylogenetic tree to plasma viral sequences sampled from the year preceding initiation of ART than to sequences from earlier in infection (20–22). This finding suggests that the replication competent latent reservoir is mainly established near the time of therapy initiation. Interestingly, Pankau *et al*. (21) tested whether a “last time point” model (HIV DNA reservoir mimics HIV RNA at the last time point prior to ART initiation) or a “cumulative” model (constant reservoir seeding and decay) better reflects observed fractions of lineage variants in the reservoir, and found that the difference between the two models was not statistically significant.

Similar contradictory findings regarding the fate of T cells in the latent reservoir have been reported. Lorenzo-Redondo *et al*. (23) and others (24) found that HIV-1 replication and evolution can occur in some tissue compartments during ART. However, studies proposing ongoing replication during ART have been criticized as being inconsistent with a vast amount of clinical data (25, 26). Several studies have found that persistence and clonal expansion of long-lived T cells, and not ongoing viral replication, was predominantly responsible for the maintenance of the latent reservoir (27–30). Cho *et al*. (31) sequenced intact proviruses from the latent reservoir, and found that the diversity decreased over time, suggesting that some sequences within the latent reservoir may be replaced as clones of infected cells expand.

Recently, White *et al*. (32) found that the CD4^+^ T cells containing intact HIV proviruses constitute at least two different subsets, each with different a decay rate: a fast-decaying T cell population with half-life of the order of days, and a second slow-decaying population with a half-life of the order of months. Similarly, Fray *et al*. (33) found that intact simian immunodeficiency virus (SIV) genomes decay with triphasic kinetics, and archival SIV variants persist, upon treatment with ART.

We postulate that the seemingly contradictory observations regarding the reservoir in the literature may be confounded by the presence of dynamically heterogeneous infected CD4^+^ T cell populations in the host. To test this hypothesis, we studied the dynamics of the divergence (the evolutionary distance from the founder sequence/s) of non-defective *env* sequences in the proviruses in CD4^+^ T cells of SIV-infected rhesus macaques on ART, and found that the divergence undergoes a biphasic decline over time. We estimated from data the distributions of proviral divergence in the short- and long-lived subsets of CD4^+^ T cells at the start of ART, and found that the short-lived cells harbor more diverged provirus from later in infection, while long-lived cells harbor less diverged provirus from earlier in infection. Furthermore, we developed a mathematical model to predict divergence kinetics during ART given these initial distributions and decay rates, and found that the model captures the observed biphasic decline of the divergence.

## Results

### Biphasic divergence decline of proviral SIV in CD4^+^ T cells during ART

Ten Indian-origin rhesus macaques were infected with simian immunodeficiency virus (SIVmac251 stock) and initiated ART at *≈* 48 weeks post-infection. Intact proviruses in CD4^+^ T cells were quantified using the intact proviral DNA assay (IPDA) (33, 34) periodically from the day ART was initiated (*t* = 0 weeks), for about four years (*t* = 216 weeks). Envelope gene (*env*) sequences in SIV DNA in circulating CD4^+^ T cells were then obtained using single genome sequencing, and filtered to remove any sequences with defects (deletions, premature stop codons, hypermutations, etc.) resulting in non-defective proviral *env* sequences; see Methods and Fray *et al*. (33) for details. In this work, we analyze the data for the first *≈* 3 years (*t* = 146 weeks). The overall decay dynamics are described in Fray *et al*. (33); see Methods and Supplementary Table S4 for details. Plasma viral RNA sequences were also obtained at ART initiation.

We determined the divergence of the non-defective proviral *env* sequences from the consensus sequence of the infecting virus stock. Based on the prior knowledge that at least two T cell subsets with different half-lives exist (32, 33), we tested whether the divergence decayed with time on ART, and if so, whether the decay was monophasic or biphasic. We found the decay to be biphasic with an approximately 500 times faster first phase decline than second phase decline when considering non-defective proviral DNA sequences obtained at all time points (Fig. 1) (p = 0.0019, paired twosided Wilcoxon signed rank exact test; Supplementary Table S5). The rapid decline phase lasted *≈* 11.6 weeks (range 4-28 weeks). Macaque T624 is an exception, and was found to have spontaneously controlled viral replication to low levels before ART (33). Interestingly, computing divergence of plasma viral RNA sequences at the start of ART (instead of proviral DNA sequences, keeping the reference unchanged), and integrated provirus at subsequent time points, revealed a much faster first phase divergence decline, approximately 900 times faster in the first phase than the second phase (p = 0.0039, paired two-sided Wilcoxon signed rank exact test); Fig. 1 and Supplementary Table S6. This likely reflects that fact that the pre-ART plasma virus sequences represent actively replicating virus while proviral sequences obtained at the same time point include actively replicating virus as well as archived sequences (33).

**Fig. 1.**
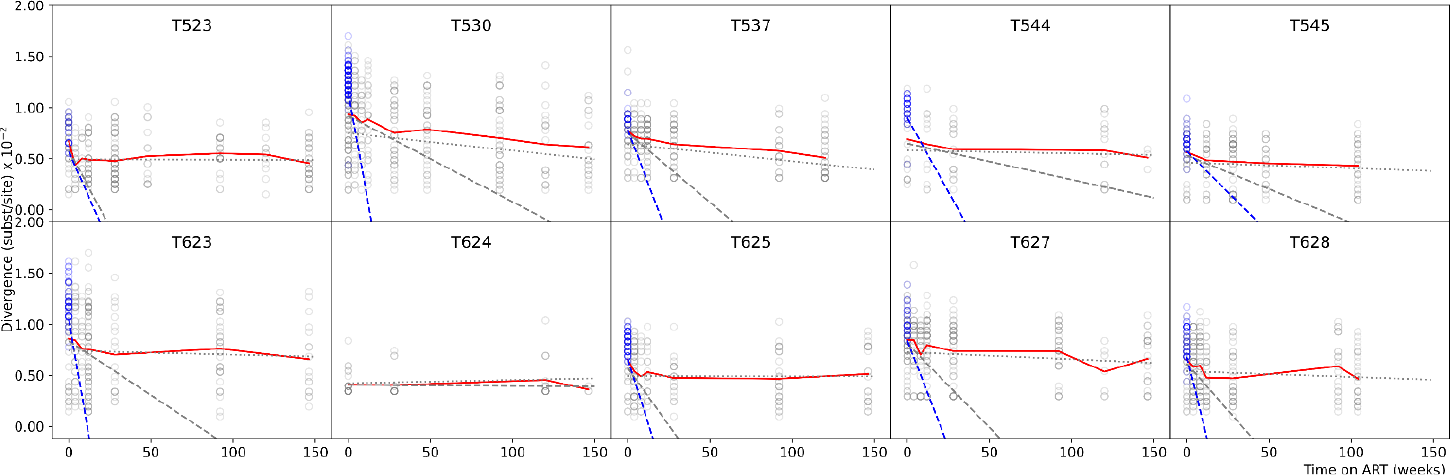
Biphasic divergence decline of non-defective provirus *env* sequences in 10 macaques infected with SIVmac251 stock, sampled longitudinally during 3 years on ART (mean divergence curve shown in red), with an initial rapid decline followed by a second slower decline. Divergence in individual experimental sequences shown in circles (non-defective proviral divergence in grey, plasma viral RNA divergence in blue). The first slope is shown by a dashed line, using non-defective proviral DNA at the start of ART in grey, and using plasma viral RNA at the start of ART in blue. The second slope is shown by a grey dash-dot line. Divergence was computed from the consensus sequence of MAC.US.x.239.M3. Plasma viral RNA could not be collected for T624.

These results show that SIV divergence of proviruses in circulating CD4^+^ T cells decline first rapidly, then more slowly over time after ART has been initiated.

### Short- and long-lived CD4^+^ T cell subsets contain provirus with different divergences

Because the non-defective provirus sequences provide the total distribution of mutations at each sampling time under suppressive ART, we separated the sequence data into sequences from the short- and long-lived cells (32, 33) by subtracting the distribution of the number of mutations measured at the last sampled time point, at which time the short-lived subset has died out, from the total distribution at the start of ART, i.e., first time point (see Methods). This showed that at the start of ART, the short- and long-lived cell subsets had different mutational distributions (Fig. 2): short-lived cells, on average, had more diverged proviral sequences than long-lived ones (p = 0.002 when non-defective proviral DNA sequences were used at all time points, and p = 0.004 when plasma viral RNA sequences were used at the start of ART; paired Wilcoxon signed rank test; Supplementary Tables S2 and S3). During untreated infection, SIV undergo continuous replication, with mutations accumulating as infection progresses (19, 37). Thus, actively replicating virus diverges further and further away from the founder virus as infection progresses (19, 38, 39), allowing the use of divergence as an indicator of the time in infection when the virus was actively replicating. Therefore, the higher divergence found in the provirus integrated in short-lived cells suggests that these viruses were deposited later in infection, closer to the start of ART. Conversely, the lower divergence found in the provirus integrated in long-lived cells indicates that these viruses were deposited earlier in infection.

**Fig. 2.**
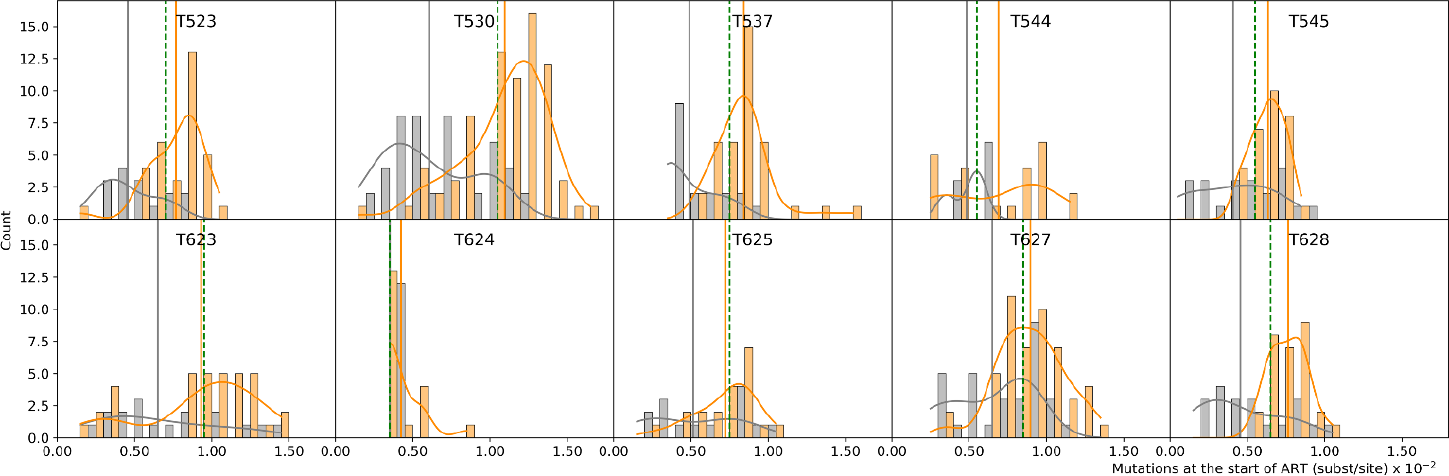
Mutational profiles of short- (orange) and long-lived (grey) populations at the start of ART, in 10 macaques infected with SIV sampled longitudinally during 3 years on ART. Orange and grey curves show the smoothed-out histograms. Solid vertical lines correspond to the mean divergence of short- (orange) and long-lived (grey) populations. Vertical green dashed lines show the median divergence of the total population.

Interestingly, the distributions of the number of mutations in the short- and long-lived cells were shifted relative to the overall divergence median - shifted right (dominated by sequences with a larger number of mutations) in the shortlived cells, and shifted left (dominated by sequences with fewer mutations) in the long-lived cells. This lends further support to the finding that the CD4^+^ T cell subsets encode different eras of virus deposition (p = 4.4e-5, Wilcoxon rank sum test); Fig 2. Again, comparing the subsequently sampled proviral sequences to the SIV RNA population at the start of ART showed that the RNA population was even more diverged, and therefore corresponds to the most recent virus variants (p = 0.0004; Wilcoxon rank sum test; Fig S6 and Supplementary Table S3).

### Decay of short- and long-lived CD4^+^ T cells with different divergences explains the biphasic divergence decline

To evaluate the observed biphasic divergence declines in SIV under ART, we developed a model to simulate the decay of provirus in two CD4^+^ T cell subsets with different decay rates and divergence distributions at the start of ART (see Methods). The simulations start with populations of cells with the estimated distributions of mutations in the short- and long-lived cell populations based on the individual experimental sequence data from each macaque. These cells then decay stochastically over time at the estimated decay rate (Supplementary Table S1). We also sample each macaque at the same times and obtain the same number of sequences as in the empirical data.

We found that the overall divergence decline predicted by the model matched the empirical decline trends; Fig. 3. We observed wide variation across macaques in the empirically estimated time for which the first phase of divergence decline lasted (bold red lines in Fig. 3), ranging from 4 to 28 weeks (average 11.6 weeks), suggesting variations in the half-lives of CD4^+^ T cells. We attempted to capture this individual heterogeneity in two ways: (i) by estimating half-lives using pooled data for all 10 macaques and then adjusting for individual variations (referred to here as mixed-effects parameters) (blue lines in Fig. 3, bold blue line showing the average of 10^3^ simulations), and (ii) by estimating half-lives separately for each macaque using only its own data (green lines in Fig. 3, bold green line showing the average of 10^3^ simulations). We found that either the mixed-effects parameters (*e*.*g*., T523, T625) or the individually estimated halflives (*e*.*g*., T628, T627, T545, T544) captured the empirically observed heterogeneity across macaques. The variability of the empirical data was based on about 20 sequences per time point (Fig 1 and Supplementary Table S4). Encouragingly, with the same sampling schedule (times and number of sequences), our simulations showed similar variability in individual simulations, matching the observed dynamics (*e*.*g*., T623, T544, T627). Similar agreement is seen between the experimental observations and model simulations when the proviral DNA sequences are replaced with plasma viral RNA sequences at the start of ART, retaining non-defective proviral DNA sequences for subsequent time points; Supplementary Fig. S7.

**Fig. 3.**
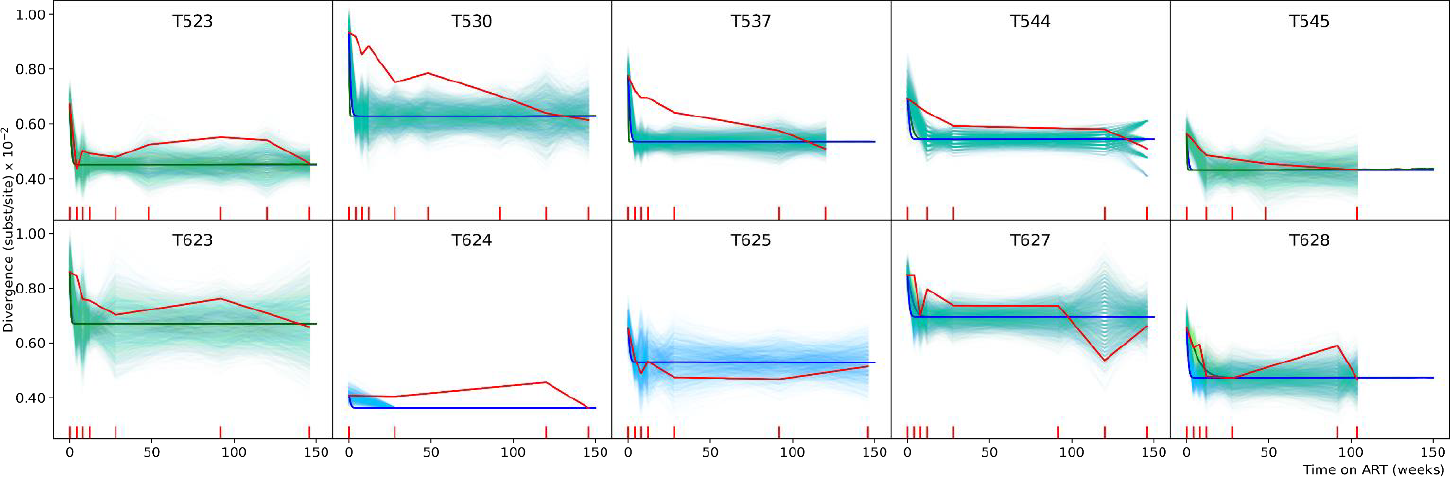
Empirically observed mean divergence dynamics (red lines) compared to model simulated divergence in 10 macaques infected with SIV sampled longitudinally during 3 years on ART. Divergence dynamics using half-lives estimated by first pooling the data of all 10 macaques, and then adjusting for individual variations are shown by blue lines. Divergence dynamics using half-lives learnt separately for each individual macaque are shown by green lines. Bold blue/green lines show the means of 103 simulations. Thin blue/green lines show individual stochastic simulations of divergence obtained by sampling according to experimental sampling times and number of sequences. Turquoise is the result of the thin blue and green lines overlapping. For macaques T624 and T625, green lines are absent as the data was insufficient to estimate half-lives individually. In T624 the thin blue lines overlap exactly with the thick blue line after *≈* 20 weeks. The red tick marks on the *x*-axis show the times when experimental samples were obtained.

### Relative contribution of parameters

We investigated the relative contribution of the model parameters to the divergence over time. Here the model parameters are: the initial frequency of short-lived cells (*f*(0)), the half-lives of short- and long-lived cells (*τ*_*S*_ and *τ*_*L*_, respectively), and the mean SIV divergence in short- and long-lived cells (*dS* and *dL*, respectively); Supplementary Methods, Figs. S8 and S9. At the start of ART, the mean SIV divergences, *dS* and *dL*, both contribute about 50%, and *f*(0) contributes 10 *-* 20% to the divergence. As time on ART proceeds, *f*(0) and *dS* become less important, while *dL* becomes dominant and eventually the sole determinant of the divergence.

### Biphasic divergence is independent of the mutational profiles in short- and long-lived CD4^+^ T cells

To investigate whether the shape of the mutation distributions of diverged provirus in short- and long-lived cell subsets has any impact on the biphasic decline, we simulated different combinations of mutation profiles (Supplementary Figs. S1 to S5). Overall, as long as the mean divergence in the short-lived cells was greater than that in the long-lived ones, qualitatively, we found biphasic divergence decline. Thus, the shape of the distribution of mutations had no qualitative impact on the overall dynamics of SIV divergence under ART. But we observed that the shape of the distribution did impact the quantitative aspects of the biphasic decline. In addition, it is possible that differences in the mutational profiles in the two cell subsets may impact other aspects of SIV within-host dynamics.

## Discussion

HIV persists as latent provirus in a reservoir of resting CD4^+^ T cells, even during suppressive antiretroviral therapy (3–5). This latent reservoir can serve as a source of viral rebound in the case of treatment interruption, making this a major hurdle to curing HIV infection (6, 7). Several seemingly contradictory views exist on when the latent reservoir is formed and what role it may have in maintaining infection, leading to different authors recommending different strategies for its elimination (10– 13, 16, 17, 19–22). In this study, we observed a biphasic decline in the divergence of non-defective proviruses in the CD4^+^ T cells of SIV infected rhesus macaques on ART. Building on the recent finding that there exist at least two subsets of CD4^+^ T cells with intact proviruses having vastly differing decay rates (32, 33), we estimated from data the divergence of proviruses harbored in short- and long-lived CD4^+^ T cells at the start of ART. We found that more diverged proviruses were harbored in short-lived cells, while less diverged proviruses were in long-lived cells. During infection, actively replicating SIV accumulate mutations and diverge further away from the founder virus as infection progresses (19, 37–39). Thus, the divergence of proviruses can serve as a proxy for the time during infection when the virus was deposited in the CD4^+^ T cell. Thus, our observations lead to the finding that SIV proviruses from later in infection (near start of therapy) are harbored in short-lived CD4^+^ T cells, while proviruses from earlier in infection are harbored in long-lived cells. We further found that in the SIV-infected macaques studied here, the mutational profile in the short-lived cells was shifted right, being dominated by viruses circulating near the start of therapy. Conversely, the mutational profile in long-lived cells was shifted left, being dominated by viruses from earlier in infection. The forces driving this shift in the mutational profiles in short- and long-lived cells are as of yet unclear, and require further investigation.

Using the estimated distribution of mutations and half-lives as inputs, we developed a mathematical model to predict the divergence dynamics during suppressive ART (assuming no viral replication or clonal expansion during suppressive ART). Our model predicts that when these cell populations decay during ART, the overall virus divergence declines in a biphasic manner, similar to that observed in our experiments. By exploring different combinations of mutational profiles at the start of ART using simulations, we found that biphasic decline occurred whenever the short-lived population had on average more diverged virus than the long-lived population.

Our model does not directly incorporate clonal expansion. However, we found that the transition from the first rapid decline in divergence to the second slower decline takes place in a matter of months (*≈*1 to *≈*7 months) in this cohort of SIV-infected macaques, well before the impact of clonal expansion becomes prominent (*≈*5 to 10 years or shorter in SIV-infected macaques on ART (33, 41, 42)). We restricted our analysis to the first *≈*3 years (146 weeks). This helped reduce the impact of a third stable CD4^+^ T cell subset with no decay observed in this cohort (33), possibly maintained through clonal expansion. Importantly, given the distributions of mutations at the start of ART, the predictions of a simple decay model can reproduce the observed divergence dynamics. This suggests that the composition of proviruses in the CD4^+^ T cell subsets at the start of therapy and the corresponding decay rates play a greater role in determining the dynamics of divergence than clonal expansion, at least over the time scales considered here.

Our experimental data, and therefore our model, investigated CD4^+^ T cells with integrated non-defective provirus *env* sequences, which may include both productively infected and latently infected cells. Thus, the predictions made here are not directly comparable to studies that focus exclusively on the latent reservoir.

Interestingly, since short-lived cells form a greater fraction of the total CD4^+^ T cell population before ART (32, 33), the overall distribution of mutations of the two subsets combined would mostly reflect the highly diverged sequences present in the short-lived cells initially after the start of ART. This agrees with recent studies showing that the majority of replication competent proviruses are genetically similar to the viruses circulating near the time of therapy initiation (20–22). As these short-lived cells die out, the overall divergence in CD4^+^ T cells increasingly reflects that of the proviruses harbored in the long-lived subset, which we found to be less diverged on average. This phenomenon explains the observed biphasic decline in overall divergence of non-defective integrated provirus. A consequence of this is that different sampling times (with respect to the start of ART) can lead to very different conclusions of how diverged the proviruses in an individual are with respect to a founder virus.

A heterogeneity has been observed in recent studies (e.g., Abrahams *et al*. (20)), where a subset of persons living with HIV (PLWH) sampled several years after the initiation of ART have proviral sequences closely resembling viruses replicating near the time of ART initiation, while a different subset of participants have proviral sequences resembling viruses replicating throughout untreated infection. We see from the macaques studied here that the duration for which short-lived cells survive can be heterogeneous (indicated by the elbow point at which the divergence decline curve changes slope, as well as a heterogeneity in half-life shown in Supplementary Table S1). This duration is also affected by phenomena such as clonal expansion. Our results suggest a possible explanation for the observed heterogeneity, indicating the possible persistence of a large fraction of short-lived cells with highly diverged proviruses in the first subset of PLWH. In the second subset, a smaller fraction of surviving short-lived cells at the time of sampling could result in a more uniform sampling of sequences with different degrees of divergence.

Thus, our results present a unifying model, suggesting that the seemingly conflicting observations in previous studies, as well as the heterogeneity observed in recent studies, may not be in conflict. Rather, they may be a result of different snapshots of the CD4^+^ T cell subsets, with variations arising from differences in sampling times, inherent stochasticity of the sampling, and individual heterogeneity.

The model proposed here has the characteristic of *last in first out* - the viruses replicating later in infection, which tend to be highly diverged, will be predominantly harbored in short-lived CD4^+^ T cells, and will be lost earlier than the long-lived cells harboring less diverged proviruses. This suggests that recent clinical trials that focus almost exclusively on viruses circulating near the start of ART (43) may miss the less diverged proviruses in long-lived cells. The good news is that people who have been on suppressive ART long enough for their short-lived cells to have died out, would tend to have less evolved proviruses which upon activation may be more susceptible to targeting by the immune system.

## Materials and Methods

### SIV data

Animals were housed at Bioqual Inc., Rockville, MD. Approval for all animal work was granted by the Institutional Care and Use Committees of Bioqual and the National Institutes of Health, and has been determined to be in accordance with the guidelines outlined by the Animal Welfare Act and Regulation (USDA) and The Guide for the Care & Use of Laboratory Animals, 8^*th*^ Edition (NIH). The regulatory standards of the indicated committees have also been followed in all the experiments which involved laboratory animals.

Ten outbred Indian-origin rhesus macaques (*Macaca mulatta*) were infected intrarectally via repetitive challenge with the SIVmac251 swarm (35) until SIV RNA was detected in the plasma using qPCR. Starting at *≈* 48 weeks postinfection, the animals were treated with a daily combination antiretroviral drug regimen of tenofovir disoproxil fumarate, emtricitabine and dolutegravir (TDF/FTC/DTG, Gilead Sciences, Inc) (33, 44). Peripheral blood mononuclear cells (PBMCs) were isolated, from which CD4^+^ T cells were purified, and DNA was extracted for quantification of infected cells using the intact proviral DNA assay (IPDA), as well as single genome sequencing of the envelope gene (33, 34, 42). Sequences with clear defects such as deletion, APOBEC3G/F-induced stop codons or hypermutation were excluded. At the start of ART (week 0), plasma viral RNA was also sequenced for 9 of 10 animals. Plasma sequences could not be recovered from animal T624 due to the low viral load at this time point.

The animals were followed for up to 216 weeks (*≈* 4 years) on ART, with longitudinal sampling of peripheral blood at weeks 0, 4, 8, 12, 28, 48, 92 104, 120, 146 and 208 or 216 post initiation of ART. When the decay kinetics of CD4^+^ T cells with intact proviruses were studied in this cohort (33), the two subsets modeled here were identified in the first < 3 years. However, between 2 and 3 years, a third stable subset with no observable decay was observed, possibly maintained by clonal expansion. In this work, we restrict our analysis to the first 146 weeks to reduce the impact of this third CD4^+^ T cell subset and clonal expansion. This dataset contains more data than the one reported in Fray *et al*. (33), with additional sequences collected from samples corresponding to weeks 0, 4, 8 and 92; Supplementary Table S4. With these additional sequences, an average of 22 sequences were obtained per animal at each time point (minimum 0, maximum 118 across animals). A complete list of the sampling times and number of sequences at each time is provided in Supplementary Table S4. Further assay details are described in (33).

The sequences reported by Fray *et al*. (33) are available in Genbank with accession numbers: OQ168980-OQ170751 (non-defective proviral DNA), and OQ168641-OQ168979 (plasma RNA) (33). Additional sequence data generated for this manuscript have been deposited in GenBank under accession numbers xx - xx.

### Divergence dynamics from data

The non-defective proviral *env* DNA sequences were aligned to a reference sequence (consensus of the SIVmac251 swarm (35)) using MAFFT (v7.490 (45)). Divergence was computed as follows

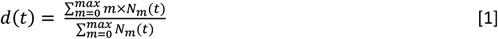

where *d*(*t*) is the divergence at time *t, m* is the raw distance (normalized by aligned sequence length) between each sequenced provirus and the reference, and *N*_*m*_(*t*) is the number of sequences at time *t* having distance *m* from the reference sequence. Considering the start of ART as time 0, *N*_*m*_(0) is estimated using experimental data as described below. *Nm*(*t*) is then computed for each time *t* by modeling the decay of cells based on their half-life. The R *lm* function (46) was used to fit a linear model to identify the slopes in the divergence curve given by the mean divergence at each time point. The elbow point at which the slope of the divergence curve changes was identified by the time point at which the absolute value of the ratio of the resulting slopes (|slope 1/slope 2|) was the largest. The slopes were estimated using at least two time points, and the time point at the potential elbow was used for both the first and second slope. These computations were carried out in R (v4.2.0 (46)), using packages ape (v5.6.2 (47)) and ips (v0.0.11 (48)).

### Estimation of half-lives

As in Fray *et al*. (33), a non-linear mixed effect approach was used to fit a biphasic decline model to non-defective proviral SIV DNA measured in the 10 macaques under ART. The biphasic decline model is given by *y* = *y*0(*fe*^*-b*1*t*^ + (1 *-f*)*e*^*-b*2*t*^), where *y* is non-defective provirus and *y*_0_ is its baseline value. The fraction *f* decays in the first (short-lived) phase with decay rate *b*_1_, while the fraction (1 *-f*) decays in the second (long-lived) phase with decay rate *b*2. The model fitting was carried out in two ways, (i) using a mixed effects approach, where all macaques were fitted simultaneously, and (ii) using the data for each macaque separately. The two methods of estimation were carried out to account for the high degree of variability across macaques. Fitting was carried out in Monolix 2021R1 (49). The estimated half-lives are provided in Supplementary Table S1.

### Estimation of initial divergence distributions

To estimate the distribution of mutations in non-defective provirus of infected cells at the beginning of ART in the short- and long-lived CD4^+^ T cells, the following assumptions were made: (i) the sequences at the start of ART come from a mixture of short- and long-lived cells, and (ii) the sequences at the last time point come only from long-lived cells, *i*.*e*., when all the short-lived cells have died out. Thus, to obtain the distribution of mutations in the long-lived cells at the start of ART (*D*_*L*_(*m*,0)), we first scale the distribution of mutations in the total set of sequences at the last measured time point (*D*_*T*_(*m,t_max_*)) to account for decay using the scaling factor 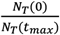. Here, *N*_T_ (0) and *N*_T_ (*t*_*max*_) refer to the total number of sequences at the start of ART and at the last sampled time point, respectively. We then multiply by (1 *-f*) to ensure that the fraction of long-lived cells at the start of ART matches the value estimated from data. The distribution of mutations in the short-lived cells at the start of ART (*D*_*S*_(*m*,0)), is then a direct subtraction of the estimated *D*_*L*_(*m*,0) from the total distribution of mutations at the start of ART (*D*_*T*_(*m*,0)). This can be written as

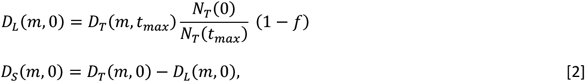

We obtained a histogram of mutations by defining each bin to represent 1 mutation. The same procedure was followed whether the data for the start of ART comprises non-defective proviral DNA or plasma viral RNA.

### Model of divergence kinetics

We developed a model of the dynamics of sequence divergence based on different initial distributions of mutations in short- and long-lived cells, and the corresponding dynamics of these cells once treatment was started, assuming no new cell infections and no cell proliferation; Fig. 4. Each of the short- and long-lived cell populations had a distribution of mutations reflecting the divergence of the sequences at the initiation of ART. The initial distributions were either the experimentally estimated distributions or idealized profiles for theoretical simulations (linear, exponential, or uniform distribution profiles). The full set of parameter values for the model are provided in Supplementary Tables S1. In each time step (= 1 day), the populations decayed according to their corresponding decay rate, irrespective of which mutational bin they belong to. The divergence of the surviving cells was computed according to Equation 1. The model was run for the same amount of time as the experiments, or for the theoretical simulations for 4 years, with 10^3^ runs for each pair of initial distributions.

**Fig. 4.**
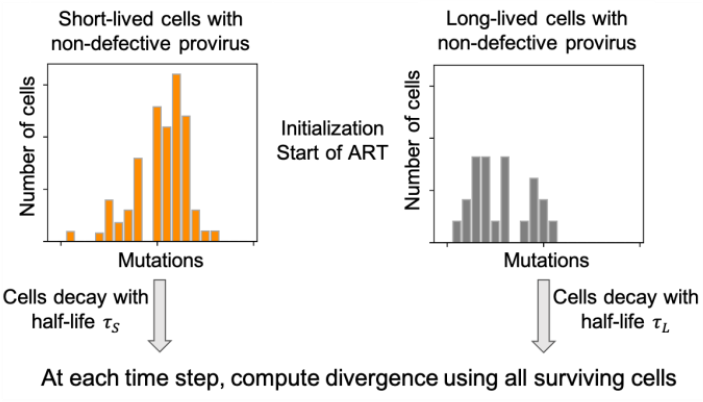
Schematic figure of the decay model used in this work. Divergence is computed using Eq. 1

Divergence was computed analytically as follows. *N*_*m*_(*t*), the number of cells with provirus having *m* mutations at any time *t*, was computed as

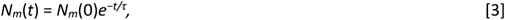

where *N*_*m*_(0) is the number of sequences with *m* mutations at time 0, and *τ* is the half-life (divided by ln(2)).

All decay simulations were carried out in Python 3.8.12 using packages pandas (v1.3.5) (50, 51) and numpy (1.20.3) (52).

## ACKNOWLEDGMENTS

Portions of this work were performed under the auspices of the U.S. Department of Energy through Los Alamos National Laboratory (LANL), which is operated by Triad National Security, LLC, for the National Nuclear Security Administration of the US Department of Energy (Contract 89233218CNA000001). Support was also provided by NIH/NIAID grants R0O1AI087520 (to T.L.), UM1 AI164561 (to R.M.R), PAVE-UM1AI164566 (to F.R.S), and R01-AI028433, R01-OD011095 (to A.S.P.). N.S. was supported by a LANL LDRD Fellowship 20210959PRD3. F.R.S was also supported by the Office of the NIH Director and National Institute of Dental & Craniofacial Research (DP5OD031834), and the Johns Hopkins University CFAR (P30AI094189). Support was also provided by the NIH Martin Delaney Collaboratory grant UM1AI164556 to D.H.B and by Howard Hughes Medical Institute to R.F.S. Animal studies were supported by AI124377, AI128751, AI149670, AI164556, AI169615 for D.H.B.

