## Supplementary Information for "Last in first out: SIV proviruses seeded later in infection are harbored in short-lived CD4^+^ T cells"

***Responsiveness of divergence to decay parameters.*** The divergence of the total population of CD4[+] T cells with intact provirus can be written in terms of the divergence of fast- and slow-decaying cells as

$$d(t) = f(t)d_F + (1 - f(t))d_S, \quad [1]$$
$$d(t) = f(t)(d_F - d_S) + d_S, \quad [2]$$

where $d(t)$ is the divergence of all CD4[+] T cells with intact provirus, $d_F$ and $d_S$ is the divergence of the fast- and slow-decaying cells, respectively, and $f(t)$ is the fraction of fast-decaying cells at any time $t$. By definition and using the exponential decay equation, $f(t)$ can be written as

$$f(t) = \frac{N_F(t)}{N_F(t) + N_S(t)}, \quad [3]$$
$$f(t) = \frac{N_F(0)e^{-t/\tau_F}}{N_F(0)e^{-t/\tau_F} + N_S(0)e^{-t/\tau_S}}, \quad [4]$$

where $\tau_F \times \log(2)$ and $\tau_S \times \log(2)$ are the half-lives of the fast- and slow-decaying cells, respectively; $N_F(t)$ and $N_S(t)$ are the number of fast- and slow-decaying cells at time $t$, where $t = 0$ corresponds to the start of ART.

The quantities that can be estimated experimentally, or derived from experimental observations, are $N_T(0)$, the total number of CD4[+] T cells with intact provirus at the start of ART, and $f(0)$, the fraction of fast-decaying cells at start of ART. That is,

$$f(0) = \frac{N_F(0)}{N_F(0) + N_S(0)} = \frac{N_F(0)}{N_T(0)}. \quad [5]$$

Therefore,

$$N_F(0) = N_T(0)f(0), \quad [6]$$
$$N_S(0) = [1 - f(0)]N_T(0). \quad [7]$$

So we rewrite equation 4 as follows

$$f(t) = \frac{f(0)e^{-t/\tau_F}}{f(0)e^{-t/\tau_F} + [1 - f(0)]e^{-t/\tau_S}}. \quad [8]$$

Substituting equation 8 into equation 2, we get

$$d(t) = \frac{d_F f(0)e^{-t/\tau_F} + d_S[1 - f(0)]e^{-t/\tau_S}}{f(0)e^{-t/\tau_F} + [1 - f(0)]e^{-t/\tau_S}}. \quad [9]$$

Thus, the divergence depends on the following parameters:

1. $f(0)$,

2. $\tau_F$,

3. $\tau_S$,

4. $d_F$, and

5. $d_S$.

We now compute the elasticity of these parameters. Elasticity estimates the effect of a proportional change in the parameter values on divergence. We further normalize the elasticities to obtain probabilities.

**Fraction of fast-decaying cells at start of ART, *f*(0):**

$$\frac{\partial d(t)}{\partial f(0)} = \frac{(d_F - d_S)e^{-t/\tau_F}e^{-t/\tau_S}}{\left(f(0)e^{-t/\tau_F} + [1 - f(0)]e^{-t/\tau_S}\right)^2}, \tag{10}$$

$$E(f(0)) = \frac{\frac{\partial d(t)}{\partial f(0)}}{\frac{d(t)}{f(0)}} = \frac{(d_F - d_S)e^{-t/\tau_F}e^{-t/\tau_S}f(0)}{\left(d_F f(0)e^{-t/\tau_F} + d_S[1 - f(0)]e^{-t/\tau_S}\right)\left(f(0)e^{-t/\tau_F} + [1 - f(0)]e^{-t/\tau_S}\right)}, \tag{11}$$

$$P(f(0)) = \frac{E(f(0))}{E(f(0)) + E(\tau_F) + E(\tau_S) + E(d_F) + E(d_S)}. \tag{12}$$

**Half-life of fast-decaying cells, $\tau_F$:**

$$\frac{\partial d(t)}{\partial \tau_F} = \frac{(d_F - d_S)f(0)[1 - f(0)]e^{-t/\tau_F}e^{-t/\tau_S}t/\tau_F^2}{\left(f(0)e^{-t/\tau_F} + [1 - f(0)]e^{-t/\tau_S}\right)^2}, \tag{13}$$

$$E(\tau_F) = \frac{\frac{\partial d(t)}{\partial \tau_F}}{\frac{d(t)}{\tau_F}} = \frac{(d_F - d_S)f(0)[1 - f(0)]e^{-t/\tau_F}e^{-t/\tau_S}t/\tau_F}{\left(d_F f(0)e^{-t/\tau_F} + d_S[1 - f(0)]e^{-t/\tau_S}\right)\left(f(0)e^{-t/\tau_F} + [1 - f(0)]e^{-t/\tau_S}\right)}, \tag{14}$$

$$P(\tau_F) = \frac{E(\tau_F)}{E(f(0)) + E(\tau_F) + E(\tau_S) + E(d_F) + E(d_S)}. \tag{15}$$

**Half-life of fast-decaying cells, $\tau_S$:**

$$\frac{\partial d(t)}{\partial \tau_S} = \frac{-(d_F - d_S)f(0)[1 - f(0)]e^{-t/\tau_F}e^{-t/\tau_S}t/\tau_S^2}{\left(f(0)e^{-t/\tau_F} + [1 - f(0)]e^{-t/\tau_S}\right)^2}, \tag{16}$$

$$E(\tau_S) = \frac{\frac{\partial d(t)}{\partial \tau_S}}{\frac{d(t)}{\tau_S}} = \frac{-(d_F - d_S)f(0)[1 - f(0)]e^{-t/\tau_S}e^{-t/\tau_S}t/\tau_F}{\left(d_F f(0)e^{-t/\tau_F} + d_S[1 - f(0)]e^{-t/\tau_S}\right)\left(f(0)e^{-t/\tau_F} + [1 - f(0)]e^{-t/\tau_S}\right)}, \tag{17}$$

$$P(\tau_S) = \frac{E(\tau_S)}{E(f(0)) + E(\tau_F) + E(\tau_S) + E(d_F) + E(d_S)}. \tag{18}$$

**Divergence of fast-decaying cells, $d_F$ :**

$$\frac{\partial d(t)}{\partial d_F} = \frac{f(0)e^{-t/\tau_F}}{f(0)e^{-t/\tau_F} + [1 - f(0)]e^{-t/\tau_S}}, \quad [19]$$

$$E(d_F) = \frac{\frac{\partial d(t)}{\partial d_F}}{\frac{d(t)}{d_F}} = \frac{d_F f(0)e^{-t/\tau_F}}{d_F f(0)e^{-t/\tau_F} + d_S[1 - f(0)]e^{-t/\tau_S}}, \quad [20]$$

$$P(d_F) = \frac{E(d_F)}{E(f(0)) + E(\tau_F) + E(\tau_S) + E(d_F) + E(d_S)}. \quad [21]$$

**Divergence of slow-decaying cells, $d_S$:**

$$\frac{\partial d(t)}{\partial d_S} = \frac{[1 - f(0)]e^{-t/\tau_S}}{f(0)e^{-t/\tau_F} + [1 - f(0)]e^{-t/\tau_S}}, \quad [22]$$

$$E(d_S) = \frac{\frac{\partial d(t)}{\partial d_S}}{\frac{d(t)}{d_S}} = \frac{d_S[1 - f(0)]e^{-t/\tau_S}}{d_F f(0)e^{-t/\tau_F} + d_S[1 - f(0)]e^{-t/\tau_S}}, \quad [23]$$

$$P(d_S) = \frac{E(d_S)}{E(f(0)) + E(\tau_F) + E(\tau_S) + E(d_F) + E(d_S)}. \quad [24]$$

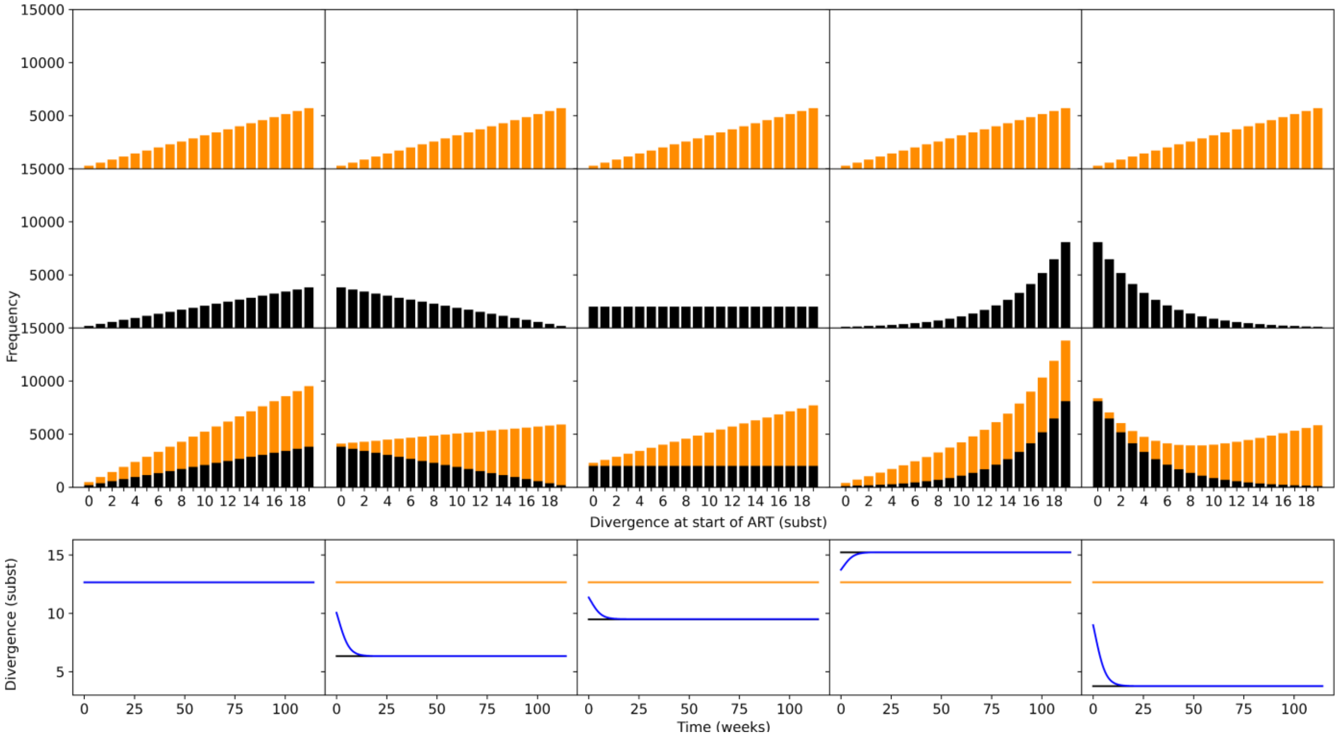


**Fig. S1.** Divergence dynamics with theoretical distributions. Here the fast-decaying population is dominated by more diverged sequences, with the divergence histogram having a linearly increasing shape. The top 3 rows show the divergence at start of ART of fast-decaying populations (row 1, orange), slow-decaying populations (row 2, black), and the total population (row 3). Row 4 shows the divergence dynamics from start of ART to ≈ 150 weeks for fast-decaying cells (orange), slow-decaying cells (black) and total population (blue). Divergence dynamics were computed analytically. The only cases when divergence declined in a biphasic manner was when the mean divergence of the fast-decaying population was higher than that of the slow-decaying population at start of ART.

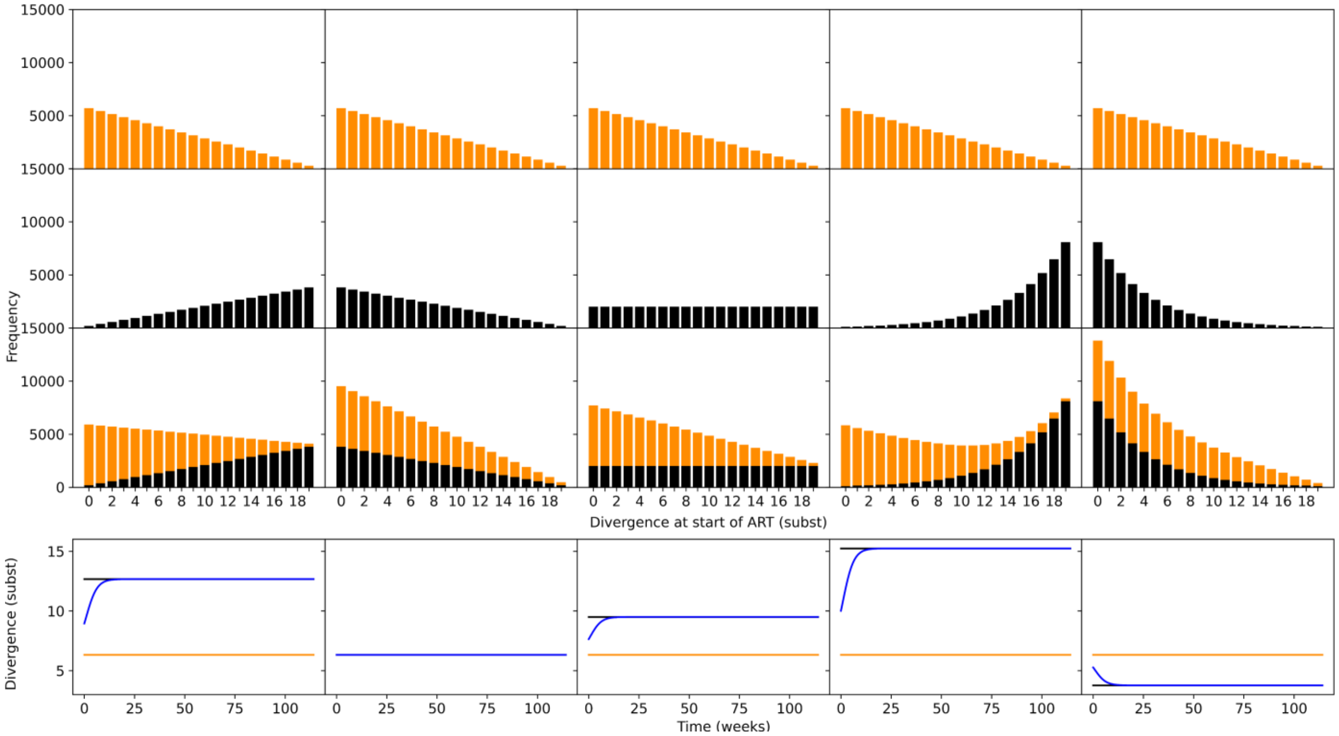


**Fig. S2.** Divergence dynamics with theoretical distributions. Here the fast-decaying population is dominated by less diverged sequences, with the divergence histogram having a linearly decreasing shape. The top 3 rows show the divergence at start of ART of fast-decaying populations (row 1, orange), slow-decaying populations (row 2, black), and the total population (row 3). Row 4 shows the divergence dynamics from start of ART to ≈ 150 weeks for fast-decaying cells (orange), slow-decaying cells (black) and total population (blue). Divergence dynamics were computed analytically. The only cases when divergence declined in a biphasic manner was when the mean divergence of the fast-decaying population was higher than that of the slow-decaying population at start of ART.

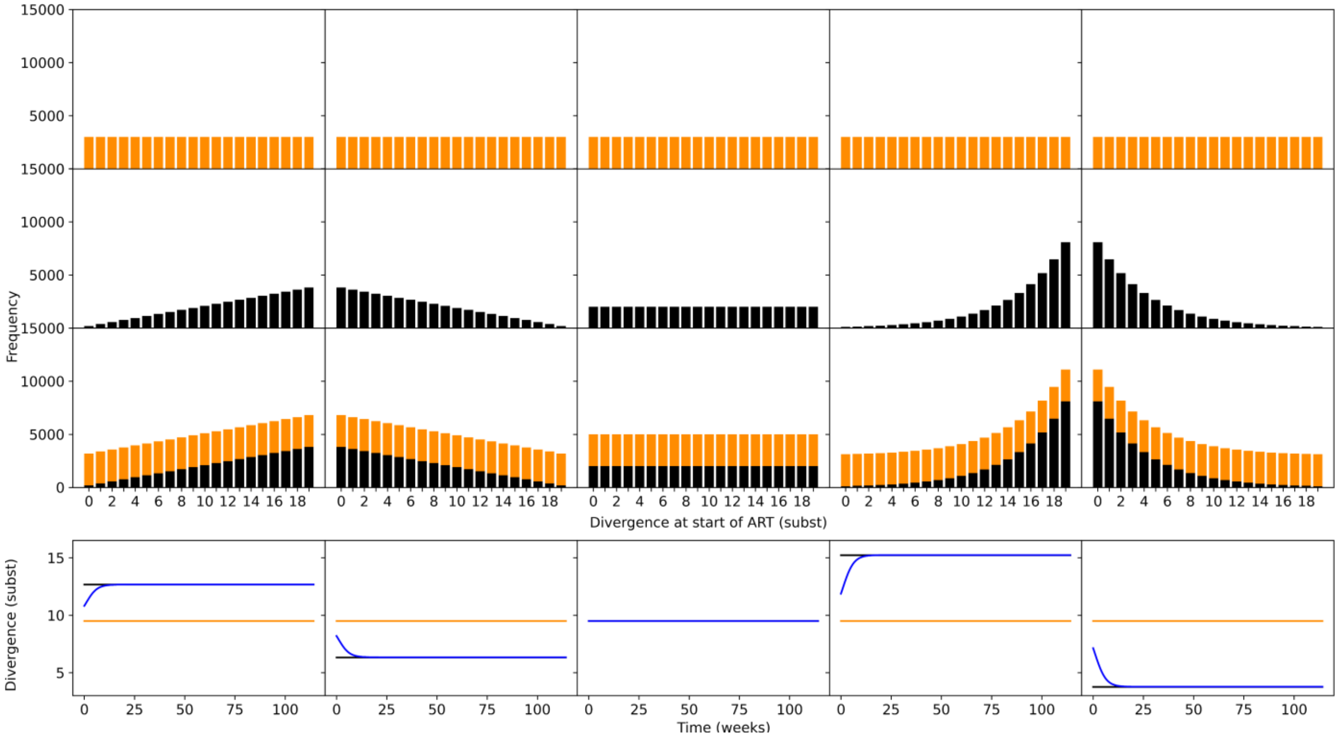


**Fig. S3.** Divergence dynamics with theoretical distributions. Here the fast-decaying population has an equal representation of different divergence values, with the divergence histogram having a uniform shape. The top 3 rows show the divergence at start of ART of fast-decaying populations (row 1, orange), slow-decaying populations (row 2, black), and the total population (row 3). Row 4 shows the divergence dynamics from start of ART to ≈ 150 weeks for fast-decaying cells (orange), slow-decaying cells (black) and total population (blue). Divergence dynamics were computed analytically. The only cases when divergence declined in a biphasic manner was when the mean divergence of the fast-decaying population was higher than that of the slow-decaying population at start of ART.

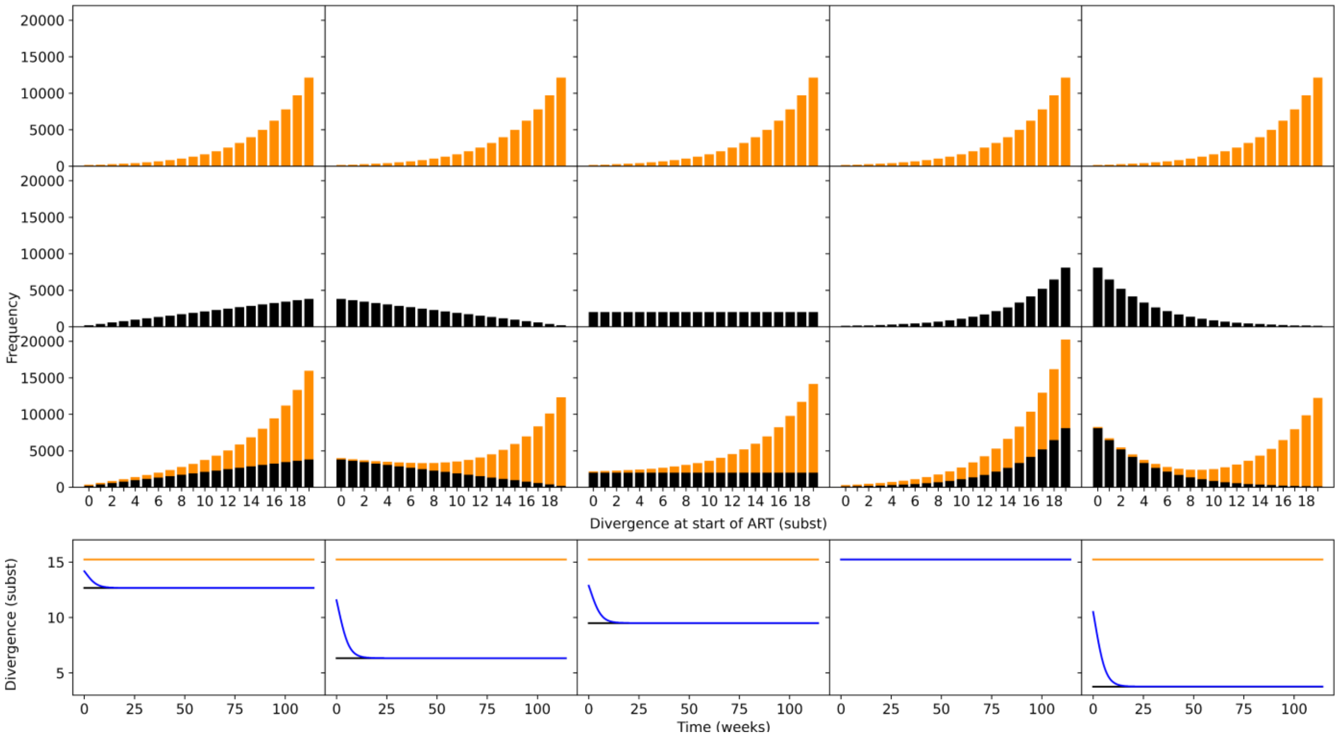


**Fig. S4.** Divergence dynamics with theoretical distributions. Here the fast-decaying population is dominated by more diverged sequences, with the divergence histogram having an exponentially increasing shape. The top 3 rows show the divergence at start of ART of fast-decaying populations (row 1, orange), slow-decaying populations (row 2, black), and the total population (row 3). Row 4 shows the divergence dynamics from start of ART to ≈ 150 weeks for fast-decaying cells (orange), slow-decaying cells (black) and total population (blue). Divergence dynamics were computed analytically. The only cases when divergence declined in a biphasic manner was when the mean divergence of the fast-decaying population was higher than that of the slow-decaying population at start of ART.

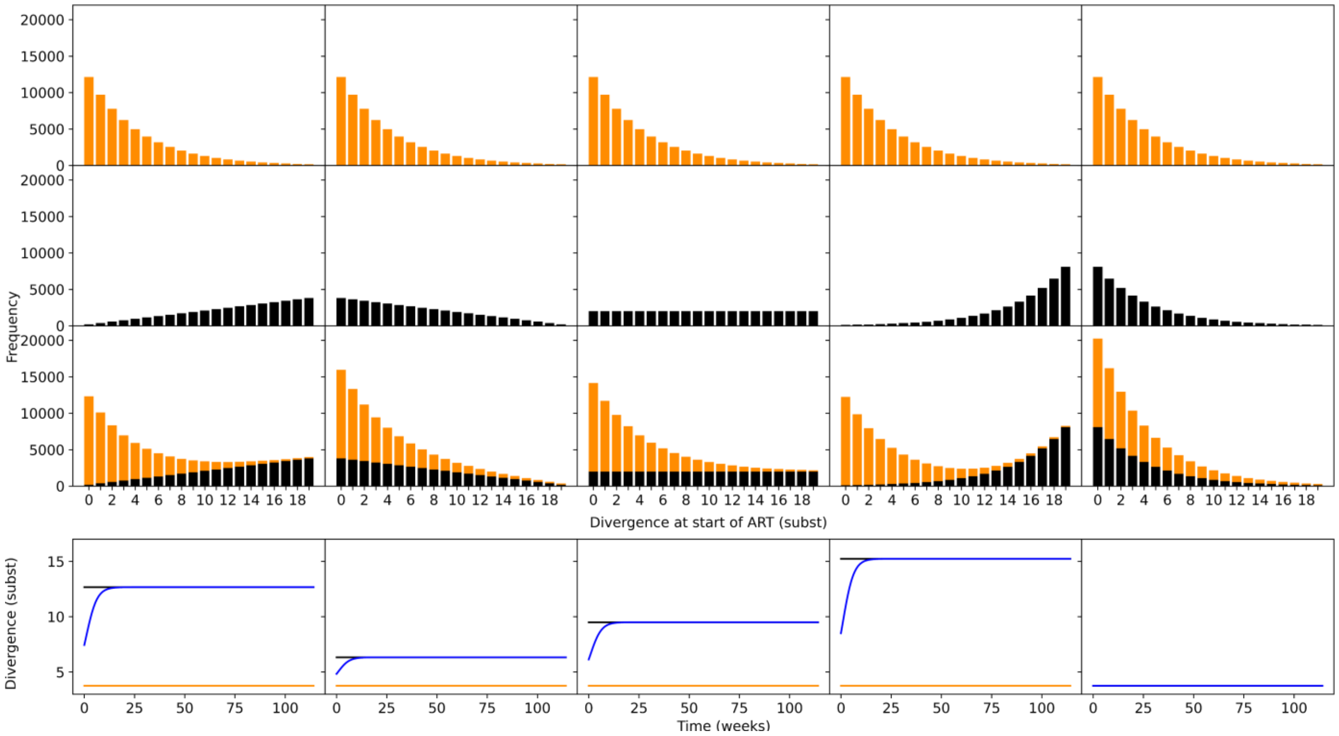


**Fig. S5.** Divergence dynamics with theoretical distributions. Here the fast-decaying population is dominated by less diverged sequences, with the divergence histogram having an exponentially decreasing shape. The top 3 rows show the divergence at start of ART of fast-decaying populations (row 1, orange), slow-decaying populations (row 2, black), and the total population (row 3). Row 4 shows the divergence dynamics from start of ART to ≈ 150 weeks for fast-decaying cells (orange), slow-decaying cells (black) and total population (blue). Divergence dynamics were computed analytically. The only cases when divergence declined in a biphasic manner was when the mean divergence of the fast-decaying population was higher than that of the slow-decaying population at start of ART.

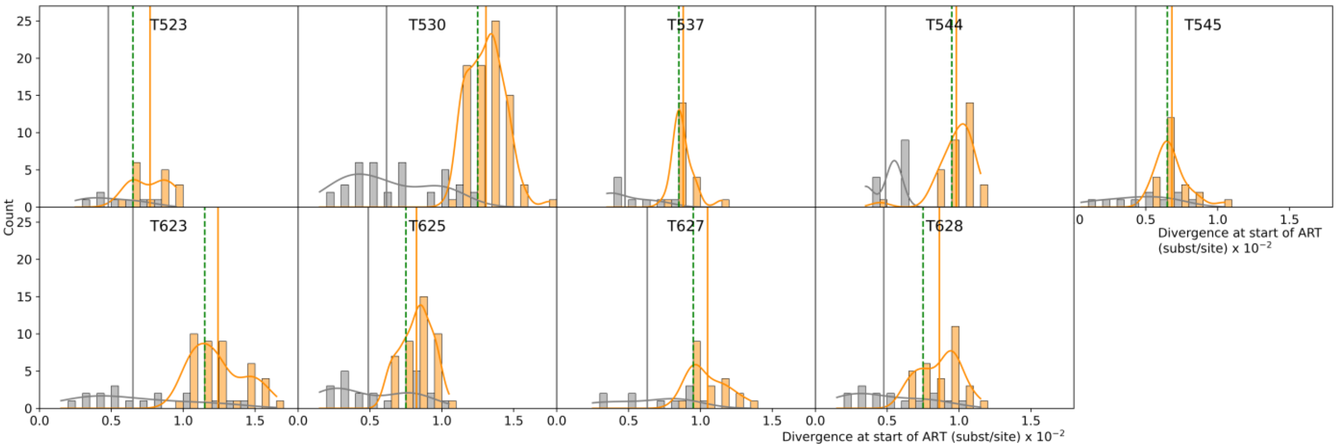


**Fig. S6.** Histograms of divergence of fast- (orange) and slow-decaying (grey) populations using plasma viral RNA sequences at start of ART in 9 SIV-infected rhesus macaques. Plasma viral RNA sequences could not be collected for macaque T624. Orange and grey curves show the smoothed-out histograms. Solid vertical lines correspond to the mean divergence of fast- (orange) and slow-decaying (grey) populations. Vertical green dashed lines show the median divergence of the total population.

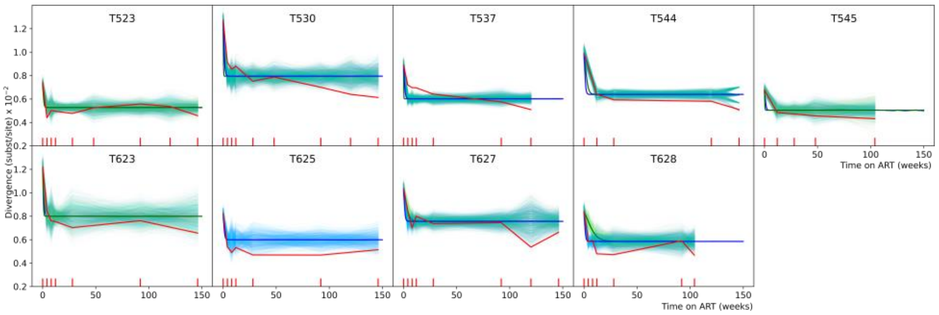


**Fig. S7.** Empirically observed mean divergence dynamics (red lines) compared to model simulated divergence. Divergence dynamics using half-lives estimated by pooling the data of all macaques are shown by blue lines. Divergence dynamics using half-lives learnt separately for each individual macaque are shown by green lines. Corresponding bold lines show means of $10^3$ simulations. Thin green/blue lines show individual stochastic simulations of divergence obtained by sampling according to experimental sampling times and number of sequences. Turquoise is the result of the blue and green lines overlapping. For macaque T625, green lines are absent as the data was insufficient to estimate half-life individually. The red tick marks on the x-axis show the times when experimental samples were obtained. The x-axis for the histograms is shown at the top of the figure, while the x-axis for the divergence dynamics plots is shown at the bottom of the figure. Simulations largely track observed change in divergence even when plasma viral RNA is used to compute divergence at start of ART.

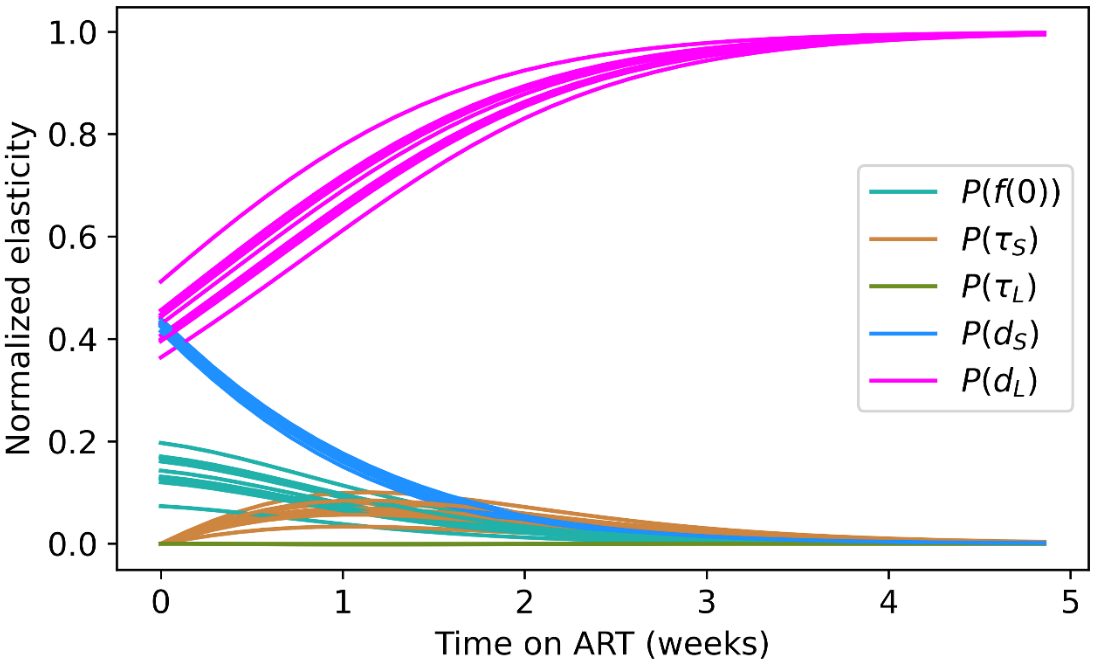


**Fig. S8.** Responsiveness to parameters was estimated by the normalized mathematical elasticity, $P$ , for each individual macaque (a) or human subject (b), shown by separate lines. $f(0)$ is the initial frequency of fast-decaying cells; $\tau_F$ and $\tau_S$ are the the half-lives of fast- and slow-decaying CD4$^+$ T cells, respectively, estimated by first pooling the data of all 10 macaques and then adjusting for individual variations; $d_F$ and $d_S$ are the initial divergence in fast- and slow-decaying cells, respectively. Details in Supplementary Methods, parameter values in Supplementary Tables S1 and S2.

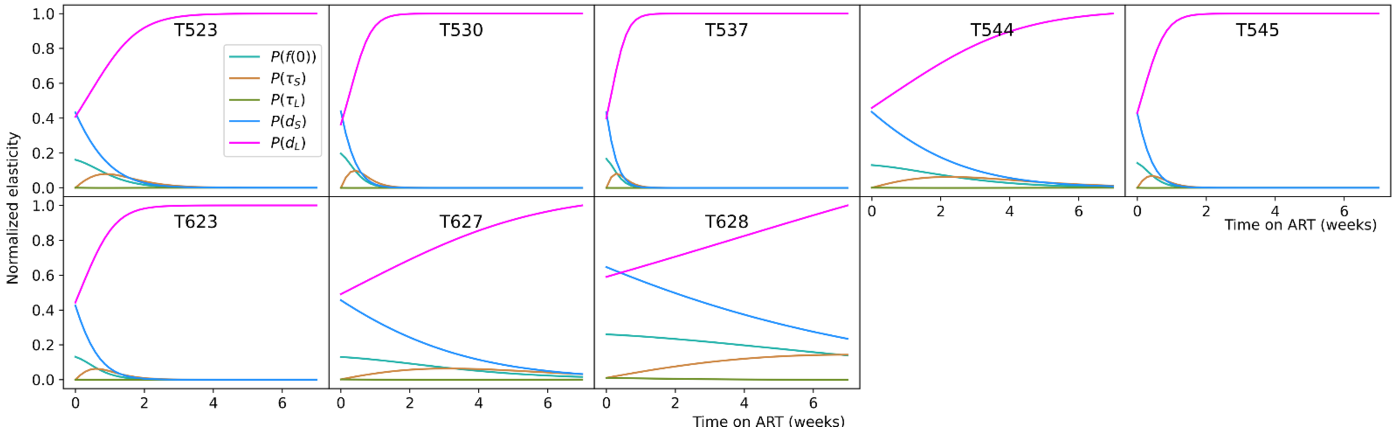


**Fig. S9.** Responsiveness to parameters was estimated by the normalized mathematical elasticity, *P* , for each individual macaque (a) or human (b), shown by separate lines. *f*(0) is the initial frequency of fast-decaying cells; $\tau_F$ and $\tau_S$ are the the half-lives of fast- and slow-decaying cells, respectively, estimated separately for each macaque (estimation did not converge for macaques T624 and T625); $d_F$ and $d_S$ are the SIV divergence in fast- and slow-decaying cells. Details in Supplementary Methods, parameter values in Supplementary Tables S1 and S2.

**Table S1. Parameter values for SIV-infected rhesus macaques.**

| Macaque ID | $d_F$ | $d_S$ | $\tau_F$ (days) mixed effects | $\tau_S$ (weeks) mixed effects | $\tau_F$ (days) individual fit | $\tau_S$ (weeks) individual fit | $f$(0) |
|---|---|---|---|---|---|---|---|
| T523 | 7.69E-03 | 4.83E-03 | 3.34 | 35 | 2.7 | 29 | 0.6 |
| T530 | 1.10E-02 | 6.07E-03 | 3.34 | 35 | 1.2 | 46 | 0.6 |
| T537 | 8.43E-03 | 5.18E-03 | 3.32 | 36 | 0.9 | 53 | 0.6 |
| T544 | 6.89E-03 | 4.83E-03 | 3.34 | 35 | 7 | 50 | 0.6 |
| T545 | 6.21E-03 | 4.15E-03 | 3.33 | 35 | 1.3 | 17 | 0.6 |
| T623 | 9.39E-03 | 6.50E-03 | 3.33 | 35 | 1.9 | 35 | 0.6 |
| T624 | 4.25E-03 | 3.50E-03 | 3.34 | 35 | - | 53 | 0.6 |
| T625 | 7.20E-03 | 5.05E-03 | 3.34 | 35 | - | 50 | 0.6 |
| T627 | 8.95E-03 | 6.42E-03 | 3.34 | 35 | 9.6 | 39 | 0.6 |
| T628 | 7.58E-03 | 4.60E-03 | 3.35 | 34 | 19.8 | 35 | 0.6 |

$d_F$ and $d_S$ are the divergence in fast- and slow-decaying cells at start of ART; $\tau_F$ mixed effects and $\tau_S$ mixed effects are the fixed effect half-lives of fast- and slow-decaying cells corrected by the random effect for each macaque, respectively, estimated by pooling the data of all 10 macaques; $\tau_F$ individual fit and $\tau_S$ individual fit are the the half-lives of fast- and slow-decaying cells, respectively, estimated separately for each macaque; $f$(0) is the frequency of fast-decaying cells at start of ART.

**Table S2. Statistics for divergence distributions at start of ART for 10 macaques using intact proviral DNA.**

| Macaque ID | Fast mean | Slow mean | p-value | Total median | Fast shift | Slow shift |
|---|---|---|---|---|---|---|
| T523 | 7.68E-03 | 4.57E-03 | 1.33E-05 | 7.00E-03 | 6.82E-04 | -2.43E-03 |
| T530 | 1.09E-02 | 6.07E-03 | 1.26E-12 | 1.05E-02 | 4.47E-04 | -4.43E-03 |
| T537 | 8.42E-03 | 4.89E-03 | 3.28E-07 | 7.50E-03 | 9.21E-04 | -2.61E-03 |
| T544 | 6.89E-03 | 4.83E-03 | 1.50E-01 | 5.50E-03 | 1.39E-03 | -6.67E-04 |
| T545 | 6.33E-03 | 4.07E-03 | 6.53E-04 | 5.50E-03 | 8.33E-04 | -1.43E-03 |
| T623 | 9.30E-03 | 6.50E-03 | 2.07E-02 | 9.50E-03 | -2.00E-04 | -3.00E-03 |
| T624 | 4.24E-03 | 3.50E-03 | 3.65E-02 | 3.50E-03 | 7.37E-04 | 8.67E-19 |
| T625 | 7.25E-03 | 5.14E-03 | 2.77E-02 | 7.50E-03 | -2.50E-04 | -2.36E-03 |
| T627 | 8.96E-03 | 6.50E-03 | 1.38E-04 | 8.50E-03 | 4.60E-04 | -2.00E-03 |
| T628 | 7.64E-03 | 4.55E-03 | 6.74E-05 | 6.50E-03 | 1.14E-03 | -1.95E-03 |

p-value reported here compares the divergence of the fast- and slow-decaying sequences per monkey at start of ART using the paired Wilcoxon exact test. The null hypothesis is the two samples come from the same distribution. A p-value $< 0.05$ indicates that the fast- and slow-decaying populations indeed have different distributions. Shift is computed as *fast (slow) shift = fast (slow) mean - total median*.

**Table S3. Statistics for divergence distributions at start of ART for 10 macaques using plasma viral RNA at timepoint 0, and intact proviral DNA thereafter.**

| Macaque ID | Fast mean | Slow mean | p-value | Total median | Fast shift | Slow shift |
|---|---|---|---|---|---|---|
| T523 | 7.69E-03 | 4.79E-03 | 3.00E-03 | 6.50E-03 | 1.19E-03 | -1.71E-03 |
| T530 | 1.31E-02 | 6.15E-03 | 3.76E-18 | 1.25E-02 | 5.78E-04 | -6.35E-03 |
| T537 | 8.80E-03 | 4.75E-03 | 1.65E-05 | 8.50E-03 | 3.00E-04 | -3.75E-03 |
| T544 | 9.81E-03 | 4.88E-03 | 3.49E-07 | 9.50E-03 | 3.13E-04 | -4.62E-03 |
| T545 | 6.82E-03 | 4.30E-03 | 2.86E-03 | 6.50E-03 | 3.18E-04 | -2.20E-03 |
| T623 | 1.24E-02 | 6.50E-03 | 1.74E-06 | 1.15E-02 | 9.27E-04 | -5.00E-03 |
| T625 | 8.24E-03 | 4.88E-03 | 8.23E-06 | 7.50E-03 | 7.38E-04 | -2.62E-03 |
| T627 | 1.05E-02 | 6.30E-03 | 3.19E-05 | 9.50E-03 | 1.00E-03 | -3.20E-03 |
| T628 | 8.63E-03 | 4.77E-03 | 2.51E-05 | 7.50E-03 | 1.13E-03 | -2.73E-03 |

p-value reported here compares the divergence of the fast- and slow-decaying sequences per monkey at start of ART using the paired Wilcoxon exact test. The null hypothesis is the two samples come from the same distribution. A p-value $< 0.05$ indicates that the fast- and slow-decaying populations indeed have different distributions. Shift is computed as *fast (slow) shift = fast (slow) mean - total median*.

**Table S4. Sampling times and number of sequences per SIV infected rhesus macaque.**

| Time (weeks) | T523 | T530 | T537 | T544 | T545 | T623 | T624 | T625 | T627 | T628 |
|---|---|---|---|---|---|---|---|---|---|---|
| 0 | 44 | 118 | 50 | 24 | 50 | 48 | 31 | 33 | 76 | 47 |
| 4 | 20 | 45 | 33 | - | - | 38 | - | 48 | 32 | 58 |
| 8 | 10 | 24 | 27 | - | - | 17 | - | 13 | 34 | 51 |
| 12 | 42 | 24 | 51 | 11 | 33 | 77 | - | 18 | 44 | 32 |
| 28 | 68 | 40 | 41 | 17 | 50 | 19 | 20 | 18 | 46 | 34 |
| 48 | 14 | 37 | - | - | 22 | - | - | - | - | - |
| 92 | 23 | 42 | 22 | - | - | 36 | - | 31 | 33 | 22 |
| 104 | - | - | - | - | 31 | - | - | - | - | 29 |
| 120 | 12 | 19 | 40 | 16 | - | - | 18 | - | 7 | - |
| 146 | 36 | 23 | - | 3 | - | 17 | 3 | 22 | 24 | - |

A '-' indicates that no sample was collected at this time.

**Table S5. Divergence slopes for 10 SIV-infected rhesus macaques on ART. Intact proviral DNA is used at all timepoints.**

| Macaque ID | Elbow point | Slope 1 | Slope 2 | \|Slope 1/Slope 2\| |
|---|---|---|---|---|
| T523 | 8 | -3.02E-04 | -2.45E-07 | 1232.05 |
| T530 | 8 | -8.42E-05 | -1.70E-05 | 4.95 |
| T537 | 4 | -1.36E-04 | -1.66E-05 | 8.19 |
| T544 | 28 | -3.54E-05 | -2.93E-06 | 12.07 |
| T545 | 12 | -6.59E-05 | -5.63E-06 | 11.69 |
| T623 | 8 | -1.06E-04 | -3.87E-06 | 27.41 |
| T624 | 28 | -9.93E-07 | 3.15E-06 | 0.32 |
| T625 | 8 | -2.40E-04 | 6.49E-08 | 3698.26 |
| T627 | 8 | -1.64E-04 | -6.97E-06 | 23.57 |
| T628 | 4 | -1.83E-04 | -5.47E-06 | 33.39 |

**Table S6. Divergence slopes for SIV-infected rhesus macaques on ART. Plasma virus RNA is used at timepoint 0, and intact proviral DNA for all subsequent timepoints.**

| Macaque ID | Elbow point | Slope 1 | Slope 2 | \|Slope 1/Slope 2\| |
|---|---|---|---|---|
| T523 | 8 | -3.64E-04 | -1.70E-07 | 2143.21 |
| T530 | 4 | -8.98E-04 | -1.95E-05 | 45.95 |
| T537 | 4 | -4.27E-04 | -1.66E-05 | 25.70 |
| T544 | 12 | -2.90E-04 | -4.87E-06 | 59.44 |
| T545 | 12 | -1.60E-04 | -5.63E-06 | 28.46 |
| T623 | 4 | -9.35E-04 | -6.59E-06 | 141.86 |
| T625 | 8 | -5.08E-04 | -8.88E-08 | 5717.00 |
| T627 | 8 | -4.16E-04 | -6.79E-06 | 61.22 |
| T628 | 4 | -6.50E-04 | -5.47E-06 | 118.71 |
